# Advanced flowering phenology of restored grasslands

**DOI:** 10.1101/2025.03.18.643883

**Authors:** Franziska Merle Willems, Johanna Bantin, Norbert Hölzel, Anna Bucharova

## Abstract

Plants introduced to degraded ecosystems in the framework of ecosystem restoration are commonly challenged by novel environmental conditions. Consequently, plant functional traits can differ between restored and reference sites, even within individual species. Studies on such intraspecific variation mainly focused on vegetative traits, while timing of life history events, phenology, received less attention so far. To address this gap, we focused on reproductive phenology of 16 flowering plant species and compared it between 47 restored meadows and 16 reference meadows in the same region. We found that plants in restored meadows flowered on average two days earlier compared to the reference, semi-natural meadows. This trend was particularly strong among early-flowering species. The potential reasons for these phenological differences are environmental differences, such as warmer microclimatic conditions and different soil properties in restored meadows; and differences in management practices, such as earlier mowing. Our findings contribute to a deeper understanding of restoration outcomes and underscore the importance of considering restoration-induced phenological shifts in conservation and management practices.

## Introduction

Ecosystem restoration is crucial to bend the curve of biodiversity decline, recover ecological functionality and stabilize the Earth’s climate (Díaz et al., 2019; IPBES, 2019; IPCC, 2019; Leclère et al., 2020). Consequently, the United Nations declared the period between 2021 and 2030 as the “Decade on Ecosystem Restoration” (United Nations General Assambly, 2019). Degraded terrestrial ecosystems often lack target vegetation, and their restoration requires reintroduction of plant diaspores from other sources (Palma & Laurance, 2015; Török et al., 2011). However, establishment and persistence of the vegetation can be challenged by the environment of restored sites.

Environmental conditions at restored sites often differ from the ones at natural sites (Brudvig et al., 2013; Klein-Raufhake et al., 2022). As restored sites are often located on former arable land that has been subject to land consolidation, there are fewer hedges, trees and other fragmenting elements (Bronstert et al., 1995; Burel & Baudry, 1990; Robinson & Sutherland, 2002). This can cause differences in microclimate, such as higher temperatures, stronger winds, and less humidity (Jacobs et al., 2022; Kanzler et al., 2019). The soils are affected by post-arable legacies, including enrichment with phosphor and lack of other essential nutrients such as nitrogen due to disrupted natural nutrient cycling (Brudvig et al., 2021; McLauchlan, 2006; Smits et al., 2008; Walker et al., 2004; Watzka et al., 2006). Land use might also differ, because restored areas are sometimes used also for other purposes, such as fodder production, timber extraction, agroforestry or infrastructure installations such as photovoltaic arrays (Bucharova et al., 2024; Dosskey et al., 2012; Silva et al., 2019; Zhang et al., 2024).

The environmental differences between restored and natural sites affect plant functional traits, which, in turn, influence ecosystem functions (Balazs et al., 2020; Pywell et al., 2003; Zirbel & Brudvig, 2020). Differences in mean community functional traits in response to environment are predominantly driven by differences in species composition (Bruelheide et al., 2018; Hodgson et al., 2011). However, evidence is emerging that there are intraspecies differences in functional traits between individuals growing at restored and natural sites (Andrade et al., 2014; Bucharova et al., 2024; Klein-Raufhake et al., 2022). Research typically focused on vegetative functional traits like plant height, leaf traits, or nutrient content in the tissues, because there is a well-established link between these traits, environment and plant performance (Chaves et al., 2002; Jung et al., 2014). Phenology-related traits received far less attention so far (but see Bucharova et al. (2024)).

Phenology, i.e. the timing of life history events such as leaf unfolding, flowering or seed set, is crucial for plant survival, growth and reproduction. Plants must synchronize their phenology with suitable environmental conditions and respond to potential changes. Most plants’ life cycles are particularly dependent on temperature, daylength and precipitation (Chmura et al., 2019). But also other factors shape plant phenology, for example nutrient content in the soil, soil water dynamics, microclimate variation, or land use (Gómez-Giráldez et al., 2020; Nomura & Kikuzawa, 2003; Nord & Lynch, 2009; Völler et al., 2013, 2017; Willems et al., 2021). Many of these factors often differ between restored and natural sites, and it is likely that this will affect plant phenology. The difference might be most apparent early in the season, as plants with early phenologies typically react stronger to environmental cues (Chmura et al., 2019).

Flowering time is a particularly important phenological trait. Apart from being critical for plant sexual reproduction, population persistence and abundance (Inouye, 2008; Wheeler et al., 2015; Willis et al., 2008), differences in flowering phenology have cascading effects at higher trophic levels through interactions with pollinators, seed herbivores and their predators community (Bucharova et al., 2016, 2022). Differences in flowering phenology between natural and restored meadows could consequently affect substantial parts of the interacting community (Bucharova et al., 2022; Franco–Cisterna et al., 2020; Johansen et al., 2019; Visser & Gienapp, 2019). Yet, such differences were rarely documented.

To address this research gap, we recorded flowering phenology of 16 flowering species that commonly grow at both restored and ancient, seminatural floodplain meadows in Germany, central Europe. The seed material for restoration was sourced from the ancient meadows and thus, population at both types of meadows had the same genetic background at the time of restoration. We hypothesize that (1) plants growing on restored meadows will flower earlier than their conspecifics on semi-natural meadows, and (2) that these differences will be greater for the earlier-flowering plants than for the later-flowering plants.

## Methods

### Study system

We studied 47 restored and 17 ancient, semi-natural floodplain meadows in the Upper Rhine Valley, Germany, scattered in an area approximately 1x3.5 km (49° 51’N, 8° 24’E). In the past, the floodplain has been covered by ancient meadows which were created by humans centuries ago for hay production and stable bedding. The long history of extensive land use has led to the establishment of species-rich plant communities, including many specialized species of high conservation value (Donath et al., 2007). A substantial part of these meadows was converted to arable land in the second half of the 20^th^ century. Between 2000 and 2007, about 70 ha have been restored back to meadow. The source of diaspores for the restoration was green hay from the remnant seminatural meadows (Hölzel et al., 2006; Hölzel & Otte, 2003). Transfer of green hay is an established restoration method for Central European meadows, which effectively introduces not only high proportion of the target species, but also the intraspecific genetic diversity of the donor populations (Dittberner et al., 2019; Hölzel & Otte, 2003; Kiehl et al., 2010). Environmental conditions at these restored sites differ from the seminatural ones. Restored sites are located on former arable land that was subject of land consolidation, while most of the seminatural meadows are located along forest edges or hedgerows (Donath et al., 2003). Soils of restored meadows are affected by agricultural legacies and often contain more phosphorus and potassium (Donath et al., 2007, 2015; Sommer et al., 2023). The land use timing also differs: restored meadows are typically mown earlier in the year, around mid-June, to preserve the nutritional value of the biomass, which is used as fodder for horses, while the remnants of ancient seminatural meadows are mown later.

### Data collection

We carried out the field work during 17 days between 27.05.2020 and 12.06.2020. We repeatedly visited each meadow every three to four days and recorded the current phenology status (see below) for 16 typical grassland species that grow at both ancient and restored sites (Table 1).

**Table 1:** List of the 16 species used in the analysis, indicating the number (N) of restored and semininatural meadows the species was growing on and the phenological status that was recorded as well as the corresponding phenology rank that represents semiquantitative gradient from early (rank 1) to late flowering (rank 12).

| Species | Restored<br>(N) | Natural<br>(N) | Phenological<br>state | Phenology<br>rank |
| --- | --- | --- | --- | --- |
| *Arabis nemorensis* | 4 | 11 | 100% Seed ripening | 5 |
| *Euphorbia esula* | 11 | 12 | 80% Seed ripening | 6 |
| *Galium album* | 49 | 12 | 5% Seed ripening | 9 |
| *Galium boreale* | 18 | 9 | 50% Flowering | 11 |
| *Galium wirtgenii* | 49 | 15 | 5% Seed ripening | 9 |
| *Genista tinctoria* | 16 | 6 | 5% Flowering | 12 |
| *Iris sibirica* | 5 | 3 | 50% Seed ripening | 8 |
| *Iris spuria* | 20 | 3 | 80% Seed ripening | 6 |
| *Leucanthemum vulgare* | 27 | 15 | 70% Seed ripening | 7 |
| *Lychnis flos-cuculi* | 16 | 9 | 5% Dispersing | 4 |
| *Medicago lupulina* | 32 | 5 | 20% Dispersing | 3 |
| *Ranunculus polyanthemos* | 6 | 11 | 70% Seed ripening | 7 |
| *Securigera varia* | 21 | 3 | 5% Flowering | 12 |
| *Tragopogon pratensis* | 39 | 6 | 50% Dispersing | 1 |
| *Trifolium campestre* | 32 | 5 | 30% Dispersing | 2 |
| *Valeriana pratensis* | 21 | 12 | 70% Flowering | 10 |

### Phenology status and rank

The selected species differed in flowering phenology and during the study period, they varied in phenology status from start of flowering to dispersing seeds. Typically, phenology studies focus on one stage, e.g. start of flowering, and record this status across time, populations and species (Chmura et al., 2019; Fitchett et al., 2015; Primack et al., 2023; Tang et al., 2016). Yet, this requires repeated visits over several months, which was logistically impossible in this study because of the global pandemic of the COVID-19 disease. We thus defined species-specific phenology status (as explained below) and recorded the day when this status was achieved at each meadow.

Specifically, we recorded the flowering phenology status that each species reached at each meadow at each visit (flowering, seed ripening or dispersing seeds) and we estimated how many % of the individuals at a given meadow reached this status. For example, when 20% of the individuals were flowering and 80% had already ripening seeds, we recorded “80% seed ripening”. After the field survey, we inspected the data and for each species, we identified the phenology status that varied most over the observation period and meadows (Table 1). For example, *Arabis nemorensis*: At the beginning of the observation period, only in a few meadows 100% individuals had ripening seeds while at the end of the period, in nearly all meadows 100% individuals had ripening seeds. The phenology status “100% seed ripening” thus best captured the variability among the meadows for this species. In other species, this phenology status was different – for example, 5% flowering in *Genista tinctoria* or 70% seed ripening in *Leucanthemum vulgare* (see Table 1). As the phenology timing for each species at each meadow, we then noted the day when this species-specific phenology status was achieved at this meadow.

To differentiate between the species along the gradient from early to late flowering, we ranked all species according to their phenological status we used in the previous analysis. The earliest flowering species had the most advanced phenology status during our field survey (*Tragopogon pratensis*, 50% dispersing seeds), and we assigned it the lowest phenology rank (= 1). On the other hand, *Genista tinctoria* and *Securigera varia* just started flowering at the time of data collection (5% flowering), and we thus assigned them the highest phenology rank (= 12). Other species were in between (Table 1). When the same phenology event was observed for two species, we assigned them the same phenology rank (see Table 1). This phenology rank thus represents a semiquantitative gradient from early to late flowering species.

### Statistical analyses

We analyzed the data in two steps. First, to understand whether the timing of flowering phenology differs between semi-natural and restored meadows, we related the phenology timing to the meadow history (semi-natural vs. restored). Second, to test whether the magnitude of the difference in flowering phenology between restored and semi-natural meadows varies between early and late flowering species, we related the species-specific effect sizes from the previous model to the species phenology rank. All analyses were carried out with the R software (R Core Team, 2023) using the *rstanarm* package (Goodrich et al., 2023).

In the first step, we tested whether phenology timing differs between restored and natural meadows. We used Bayesian multilevel regression models (also called linear “mixed models”). When phenology timing was the response variable (= y) and meadow type was the single predictor (that can take two values i, “natural” or “restored”) and y is normally distributed with variance σ, (*y_i_* ∼ Normal(*µ, σ*)) the model can be written as:

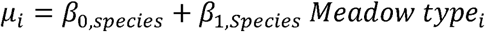

with µ_i_ = predicted phenology timing, β_0_ = intercept, which is in this case the phenology timing on natural meadows, β_1_ = the effect of meadow type, i.e. the difference in phenology timing on restored meadows. The small _species_ annotation implies that we estimated species specific deviations for the phenology timing on natural meadows (= intercept β_0_) as well as species specific effects of restoration (= β_1_). This corresponds to using species as a random factor with a random intercept and a random slope for the effect of restoration status. Thereby we estimated how much a species phenology timing differed from the average timing on natural meadows and from the average effect of restoration (i.e. the difference in timing on restored meadows compared to natural meadows). Both β_0,species_ and β_1,species_ are assumed to come from a multivariate normal distribution. For a complete model notation see supplementary material (Supporting Information Text) and for more explanations on this kind of model notation see e.g. McElreath (2018).

In the second step, we tested whether the species-specific phenology differences between the ancient semi-natural and restored meadows vary along the gradient from early and to later flowering species. To do so we extracted the estimated differences in the phenology timing between semi-natural and restored meadows (β_1,Species_) from the model described above for all 16 species. We then related these estimated differences in phenology timing as the response variable (= y) to the phenology rank as the single predictor. Given that y is normally distributed with variance σ, (*y_i_* ∼ Normal(*µ, σ*)) The corresponding model can be written as:

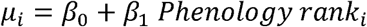

with µ_i_ = predicted average difference in phenology timing and β_1_ = the effect of phenology rank, i.e. an estimate of how much the phenological response (= difference in phenology timing) differs between early and late flowering plants.

We used the *stan_glmer* function with four MCMC chains of 2000 iterations in the *rstanarm* package (Goodrich et al., 2023; Stan Development Team, 2022) and weakly informative prior distributions (see Table S1, in the supplementary material) to reduce posterior uncertainty and stabilize the computations (Muth et al., 2018). Chain convergence and model fit and convergence was evaluated using the *shinystan* function (Gabry et al., 2022) and by ensuring that all 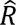 values were below 1.1., following the procedure described by Muth et al. (2018). Raw data is shown in Figure S1. For graphical and numerical checks of model fit and convergence see Table 1 and Figures S2-S6 in the supplementary material. We summarized all posterior parameter distributions with their mean and 95% probability intervals (PI), i.e. the 2.5% and 97.5% quantile of the posterior, which are the equivalent of the 95% confidence interval in a frequentist context.

## Results

Over all species, the phenology of plants on restored meadows was two days advanced in comparison to the plants growing on semi-natural meadows (µ_β1_ = -2.0, PI = [-2.8, -1.1]) (Figure 1 and Table S2). The species-specific mean differences in phenology timing varied between 2.5 days to 1.5 days (with variance σ^2^_β1_ = 0.7) and were credibly different from zero (with 95% probability) for 14 of the 16 species (Figure 1 and Tables S2 and Table S3). Overall, the estimated difference of phenology timing decreased from early flowering to late flowering plants (Figure 2, Table S3 and Table S4).

**Figure 1:**
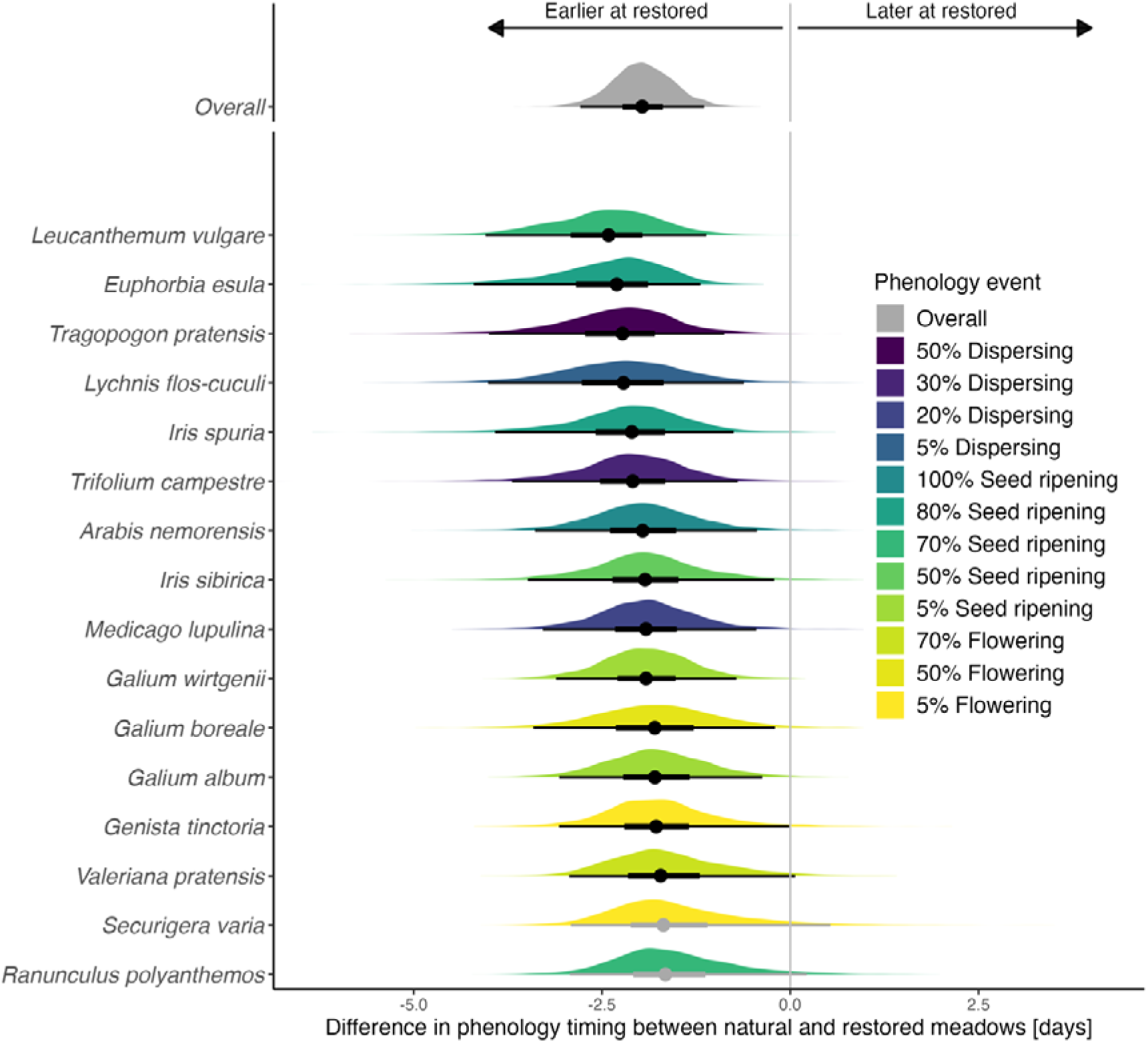
Estimated differences in phenology timing between natural and restored meadows. Negative differences indicate earlier phenology at restored meadows. Shown are posterior predictions, with points indicating the parameter estimates and whiskers their 50% (thick lines) and 95% (thin lines) probability intervals. Intervals credibly different from zero (with 95% probability) are printed in black. The average difference over all species is depicted at the top. Species are ordered by effect size (difference between the meadow types). Colors indicate the recorded phenology event for each species. (See Table 1)

**Figure 2:**
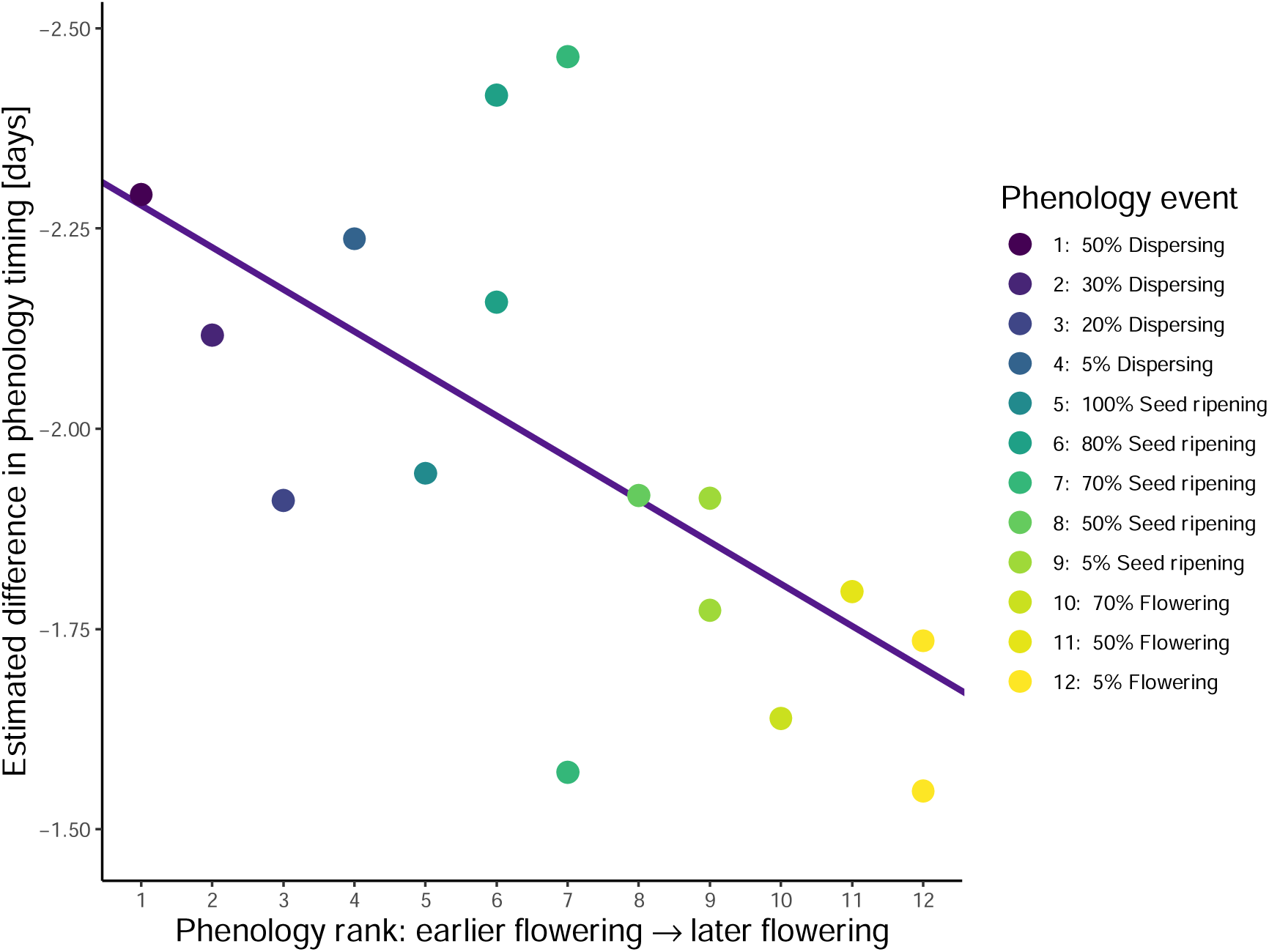
Relationship between the estimated differences in phenology timing (on restored compared to natural meadows) and phenology rank for all 16 species. Overall, the estimated differences in the phenology timing between restored and natural meadows were larger for earlier flowering plants. The purple line indicates this trend. Light purple lines are 100 draws of the fitted lines from the posterior distribution.

## Discussion

Flowering phenology is an important trait that commonly contributes to local adaptation and affects plant interactions with pollinators and seed predators (Bucharova et al., 2022; Elzinga et al., 2007; Trunschke et al., 2024; Valdés & Ehrlén, 2017). Here we show that flowering phenology of many species at restored meadows is on average two days earlier than flowering phenology of conspecifics at seminatural meadows in the same region. This trend was more pronounced in species that flowered comparatively early in the season. The mechanisms behind the phenological differences are possibly both phenotypic plasticity and adaptation to the specific conditions at seminatural and restored meadows (Anderson et al., 2012; Bucharova et al., 2024; Merilä & Hendry, 2014). In any case, differences in reproductive phenology between natural and restored meadows could have complex consequences on the interaction biota (Franco–Cisterna et al., 2020; Johansen et al., 2019; Visser & Gienapp, 2019).

Phenology differences between restored and natural meadows could be plastic reaction to microclimate, soil conditions, or local plant diversity. Many of the ancient semi-natural meadows are located along forest edges and they generally contain more trees and hedges compared to the restored meadows, which are usually located on former arable land with little structural diversity. This possibly influences local temperature and humidity, resulting in warmer conditions on restored meadows due to less shading, and consequently, earlier flowering (Piao et al., 2019; Tang et al., 2016; Willems et al., 2022). Earlier flowering at restored meadows could be also caused by soil conditions, as soils at these meadows are richer in phosphorus, and higher phosphorus supply advances flowering phenology (Donath et al., 2007; Nord & Lynch, 2008; Petraglia et al., 2014). Lastly, earlier flowering at restored meadows could be affected by lower diversity of the plant community in comparison with ancient seminatural meadows (Sommer et al 2023). Plants growing in less diverse communities have been shown to flower earlier, an effect possibly mediated through changes of microclimate and soil properties (Wolf et al 2017).

The earlier flowering at restored meadows could also reflect rapid local adaptation to management practices, specifically mowing regime (Reisch & Poschlod, 2009; Völler et al., 2013, 2017). In contrast to late mown seminatural meadows, the restored meadows are mown rather early, at the time when many of the grassland species start producing seeds. Consequently, only plants that complete their life cycle before the vegetation is mown can contribute to the next generation. In species that start seed ripening around the time when the meadows are mown, there should be rather strong selection for earlier flowering. While this process is likely, we cannot provide evidence of rapid adaptation because we observed the plants in the field and we cannot differentiate whether the observed pattern is plastic, or a result of selection. However, Bucharova et al. (2024) grew one of the species included in this study, *Galium wirtgenii*, in a common garden, and found evidence for genetic differentiation of flowering time between a seminatural and a restored meadows. This differentiation happened over just few generations, because the seed material for restoration was sourced from the studied seminatural meadow some 20 years ago, and thus, represented the same genetic pool. Such rapid evolution across just few generations has been documented also in other study systems (Conrady et al., 2023; Magnoli, 2020). Whether it is the case also in other species included, in which we detected field differences in phenology, would require a test in a common garden experiment.

The difference in phenology between restored and seminatural meadows was more pronounced in early flowering species. This is in line with previous studies (Fitter & Fitter, 2002; Freimuth et al., 2022; Kopp et al., 2020; Miller-Rushing & Inouye, 2009). A reason for this might be that variability of temperatures is greater early in the year (Menzel et al., 2006). Menzel et al. (2006) argue that, for insect pollinated plants increased temporal and spatial variability in spring phenology might make synchronization with pollinators more successful by allowing greater flexibility in the timing of both species.

Changes in plant flowering times may affect interacting species such as pollinators or seed herbivores. If the timing of phenological events shifts differently for species involved in entangled biotic interactions, this can lead to asynchrony, resulting in mismatches between species and their (food) resources (Schmitz, 2013). Disruptions in plant-pollinator interactions can result in reduced quantity and quality of pollination services (Burkle et al., 2013; Forrest, 2014; Franco–Cisterna et al., 2020). Furthermore, differences in flowering phenology may also lead to differences in seed-herbivore infestation rates (Bucharova et al., 2016). However since, the phenological differences we observed between restored and semi-natural meadows were rather modest, it is unlikely that they will cause such disruptions in plant-pollinator interactions or declines in the quantity and quality of pollination services that have been observed elsewhere (Burkle et al., 2013; Forrest, 2014; Franco–Cisterna et al., 2020). Previous research has even shown that heterogeneous flowering times at the landscape level can even improve floral resources for pollinators in agricultural landscape (Johansen et al., 2019)

In summary, we have shown, that plants growing in restored meadows differ in flowering phenology from the same species growing at seminatural sites. This adds to mounting evidence that conspecific plants growing at restored and natural sites may significantly differ in functional and life history traits (Bucharova et al., 2024; Klein-Raufhake et al., 2022).

## Author contributions

AB and NH conceived the ideas and designed methodology for this study. JB collected data. FMW led the data analysis and the writing of the manuscript. All authors contributed significantly to the drafts and gave final approval for publication.

## Supplementary material

**Figure S1:**
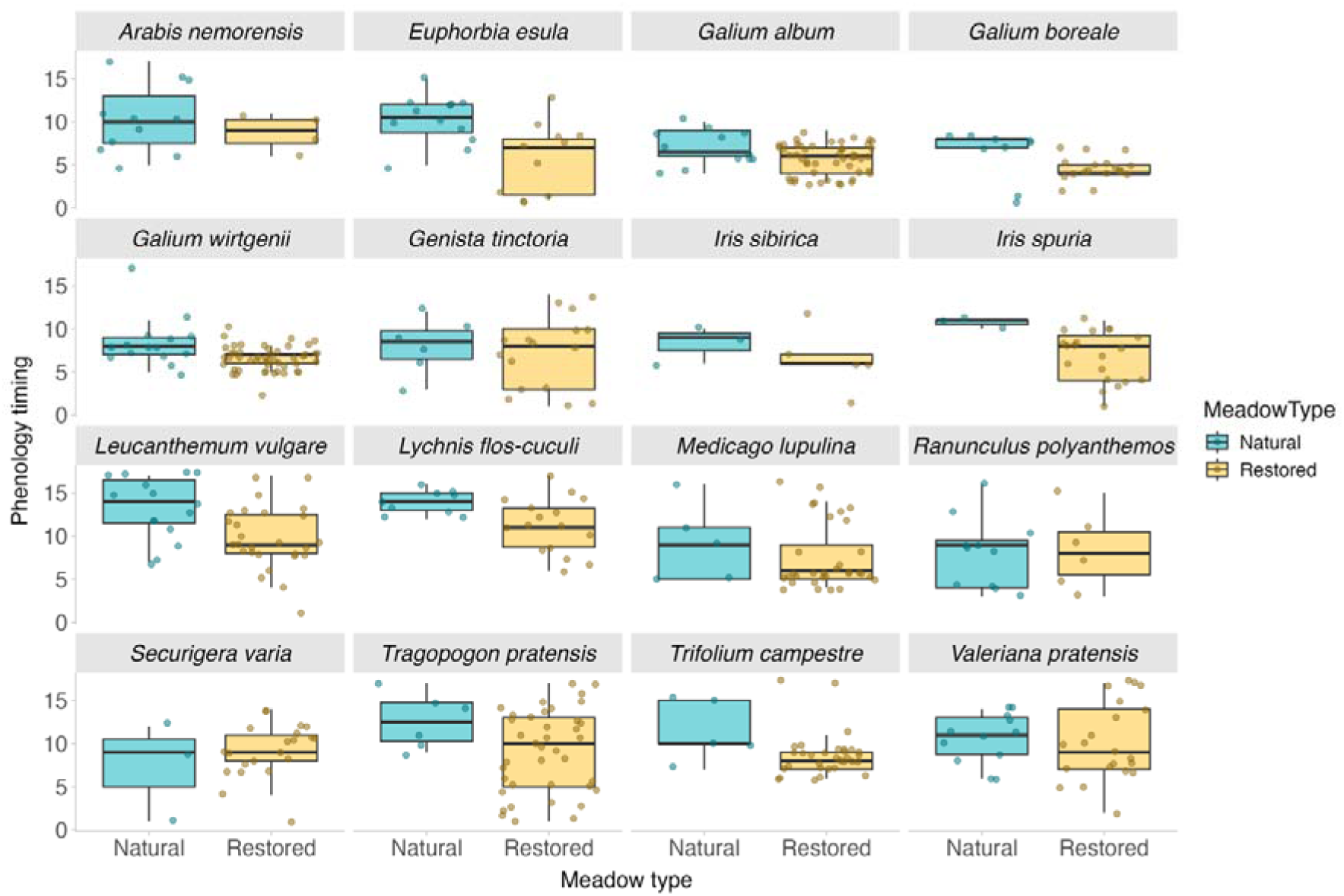
Raw data of the phenology timing for all 16 species on restored and natural meadows.

### Supporting Information Text

#### Full model notation

When phenology timing is the response variable (= y) and meadow type is the single predictor (= x, that can take two values i, “natural” or “restored”) the model can be written as:

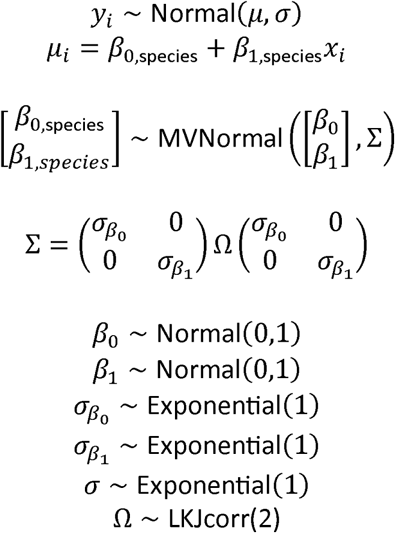

The parameters *β*_0_ and *β*_1_ are assumed to come from a multivariate normal distribution (MVKNormal). This multivariate normal distribution consists of the vector of mean parameters and a covariance matrix of standard deviations Σ. The remaining elements *β*_0_ - *σ*, are the hyper-priors; *β*_0_ - Normalc0,11 for example, refers to the population-level intercept (that is the phenology timing on natural meadows). The notation implies that the individual *β*_0,species_ come from a common distribution defined by the hyper-prior *β*_0_. Assigning a prior to *β*_0_ makes the model somewhat skeptical of individual intercepts that deviate strongly from the average. LKJcorr ist the prior for the correlation matrix. For a helpful explanation of this kind of notation see for example the excellent blog post by Will Hipson (Hipson, 2020) and McElreath’s Statistical Rethinking (McElreath, 2018).

**Table S1:**
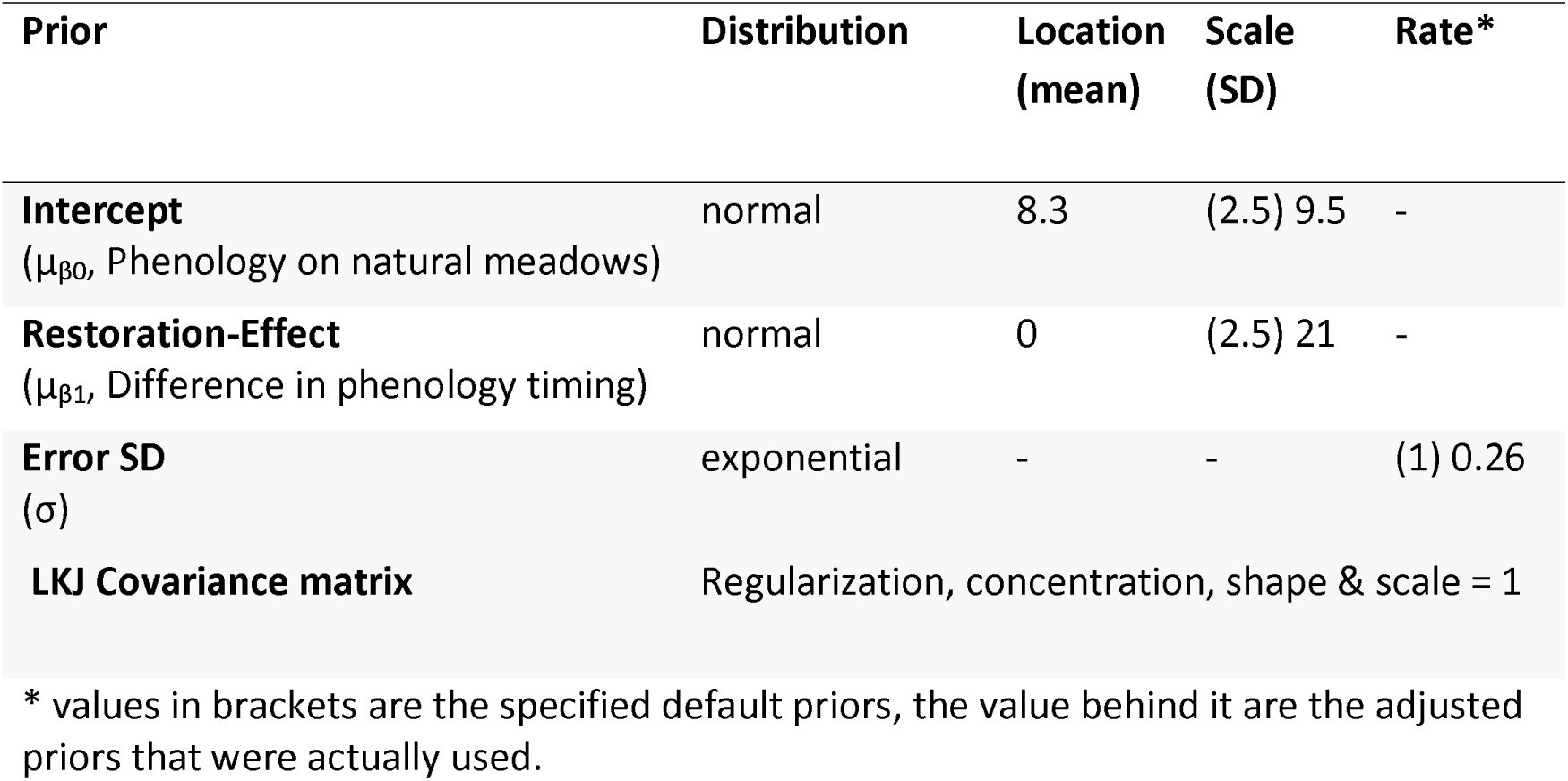
Prior distributions of the model.

| Prior | Distribution | Location<br>(mean) | Scale<br>(SD) | Rate* |
| --- | --- | --- | --- | --- |
| **Intercept**<br>($\mu_{\beta 0}$, Phenology on natural meadows) | normal | 8.3 | (2.5) 9.5 | - |
| **Restoration-Effect**<br>($\mu_{\beta 1}$, Difference in phenology timing) | normal | 0 | (2.5) 21 | - |
| **Error SD**<br>($\sigma$) | exponential | - | - | (1) 0.26 |
| **LKJ Covariance matrix** | Regularization, concentration, shape & scale = 1 |  |  |  |

\* values in brackets are the specified default priors, the value behind it are the adjusted priors that were actually used.

**Table S2:**
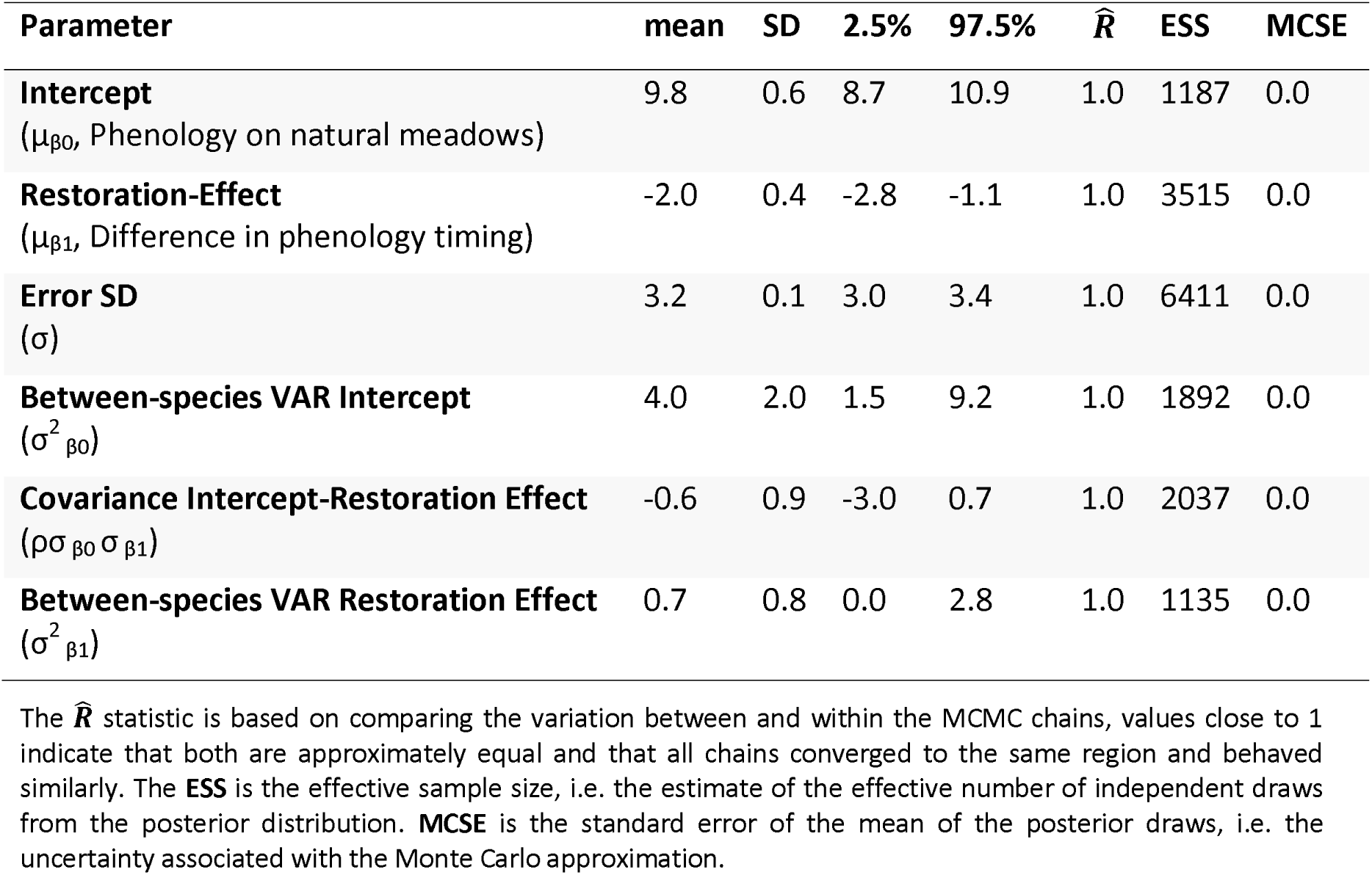
Posterior summary statistics of the estimated parameters and model diagnostics of the model investigating the effect of meadow restoration status on phenology timing. The restoration effect is the difference (in days) in the phenology timing of semi-natural meadows compared to restored meadows. Negative differences indicate that plants were flowering earlier on restored meadows.

**Table S3:**
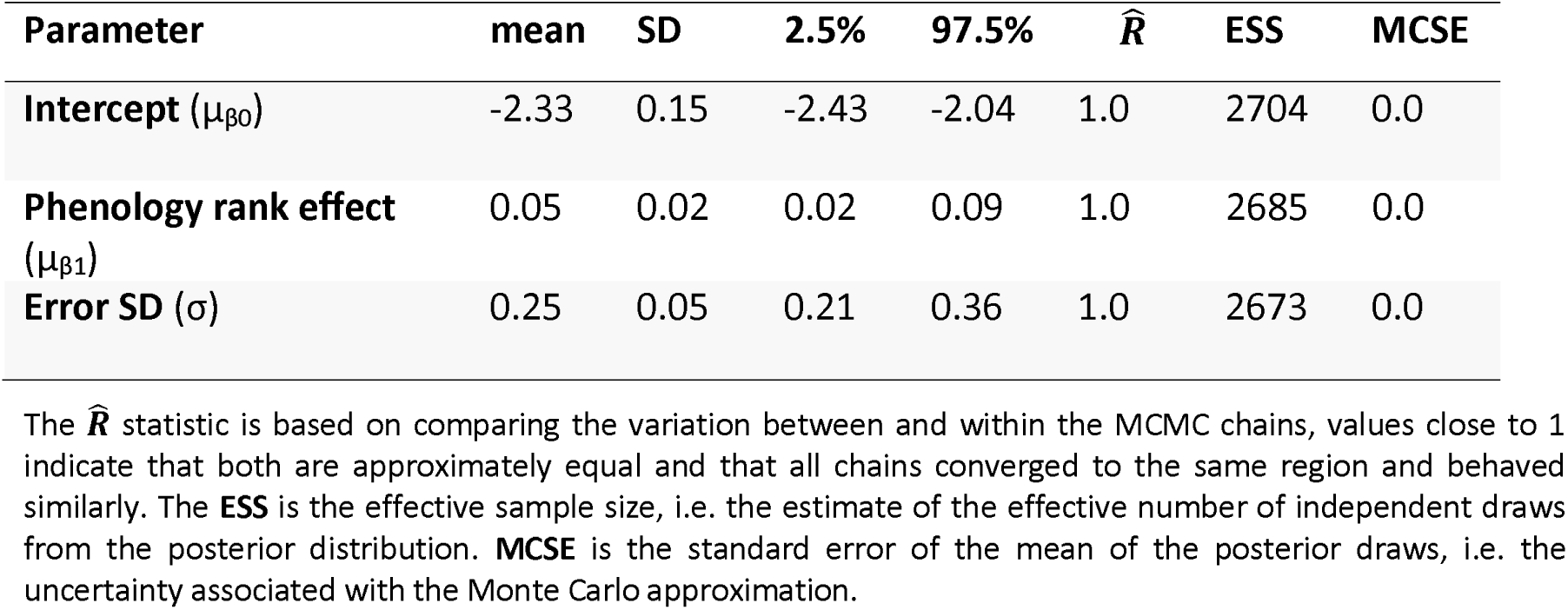
Posterior summary statistics of the estimated parameters and model diagnostics of the model investigating the relationship of differences in phenology timing (in days) on restored compared to natural meadows and phenology rank.

**Table S4:** Species specific estimates for the effect of meadows restoration status on phenology timing [differences in days]. Negative differences indicate that plants developed faster on the restored meadows compared to the natural meadows. Species are ordered by effect size, as in Figure 2. *Indicate that the 95% PI did not overlap zero.

| Species | Mean differences in<br>phenology timing | 2.5% | 97.5% |
| --- | --- | --- | --- |
| *Leucanthemum vulgare* | -2.5 * | -3.9 | -1.0 |
| *Euphorbia esula* | -2.4 * | -4.1 | -1.1 |
| *Tragopogon pratensis* | -2.3 * | -3.8 | -0.7 |
| *Lychnis flos-cuculi* | -2.2 * | -4.0 | -0.6 |
| *Iris spuria* | -2.2 * | -3.8 | -0.7 |
| *Trifolium campestre* | -2.1 * | -3.7 | -0.7 |
| *Arabis nemorensis* | -1.9 * | -3.4 | -0.4 |
| *Galium wirtgenii* | -1.9 * | -3.2 | -0.8 |
| *Iris sibirica* | -1.9 * | -3.4 | -0.1 |
| *Medicago lupulina* | -1.9 * | -3.4 | -0.5 |
| *Galium boreale* | -1.8 * | -3.5 | -0.2 |
| *Galium album* | -1.8 * | -3.1 | -0.4 |
| *Genista tinctoria* | -1.7 * | -3.2 | -0.1 |
| *Valeriana pratensis* | -1.6 * | -3.1 | -0.1 |
| *Ranunculus polyanthemos* | -1.6 | -3.0 | 0.1 |
| *Securigera varia* | -1.5 | -3.1 | 0.3 |

**Figure S2:**
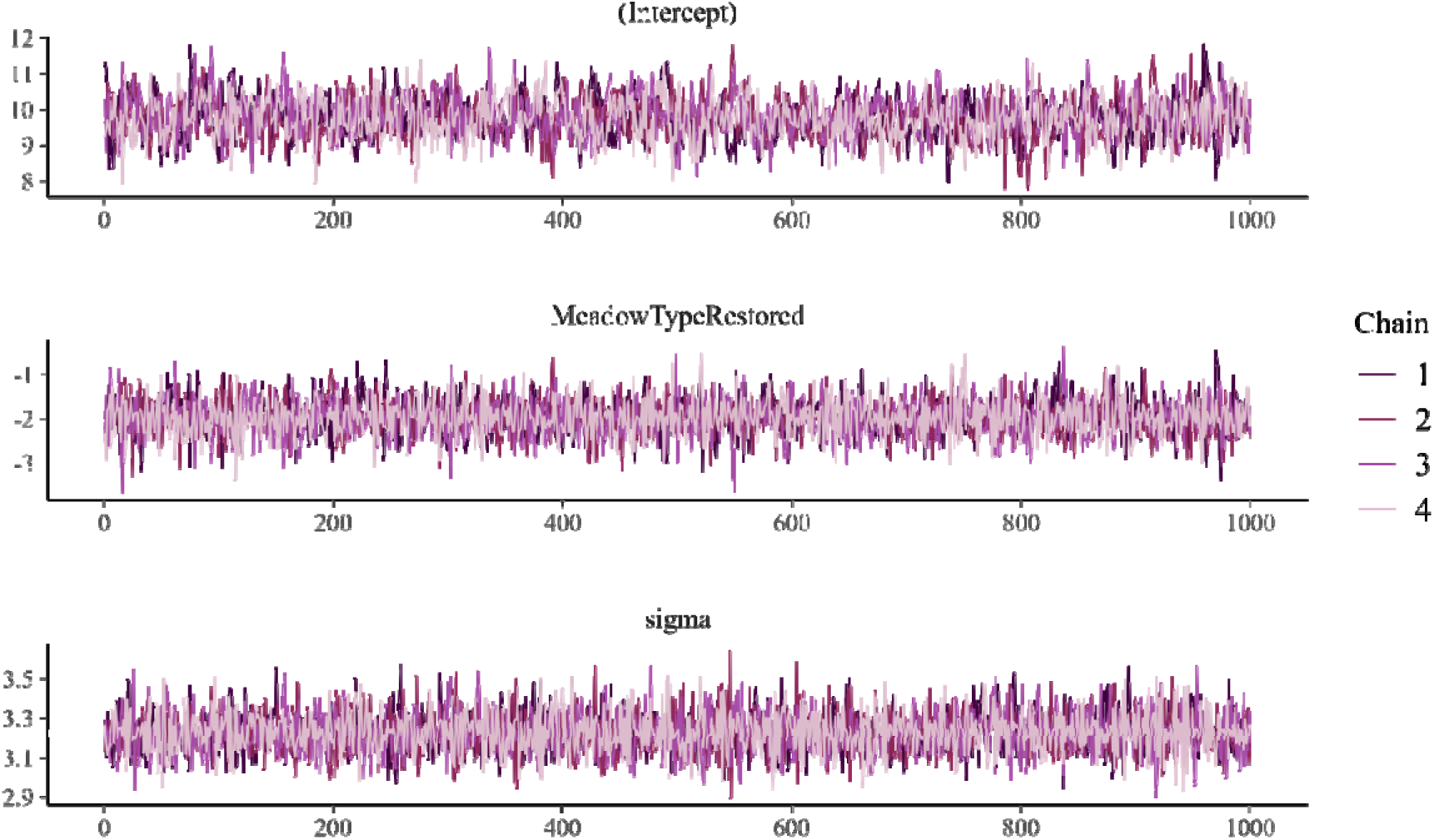
Trace plot for the Intercept (β_0_, Phenology on natural meadows), Meadow type (β_1_, Difference in phenology timing on restored meadows) and sigma. The four chains show no discernible pattern, a strong sign of model conversion.

**Figure S3:**
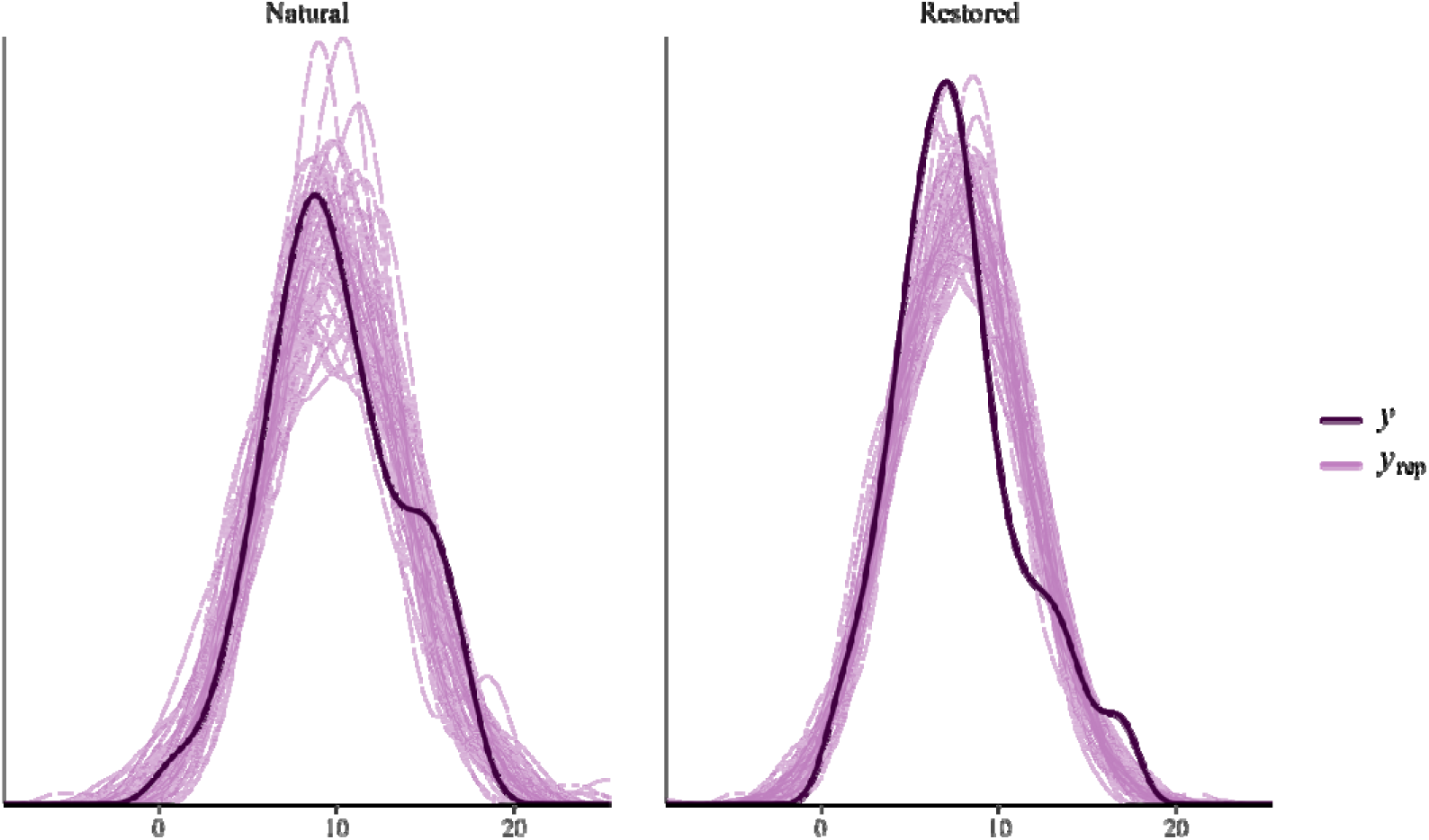
Graphical posterior predictive check comparing the observed distribution of phenology timing (y = dark purple lines) to 100 simulated datasets from the posterior predictive distribution (y_rep_ = light purple lines), for the phenology on natural meadows (left) and restored meadows (right).

**Figure S4:**
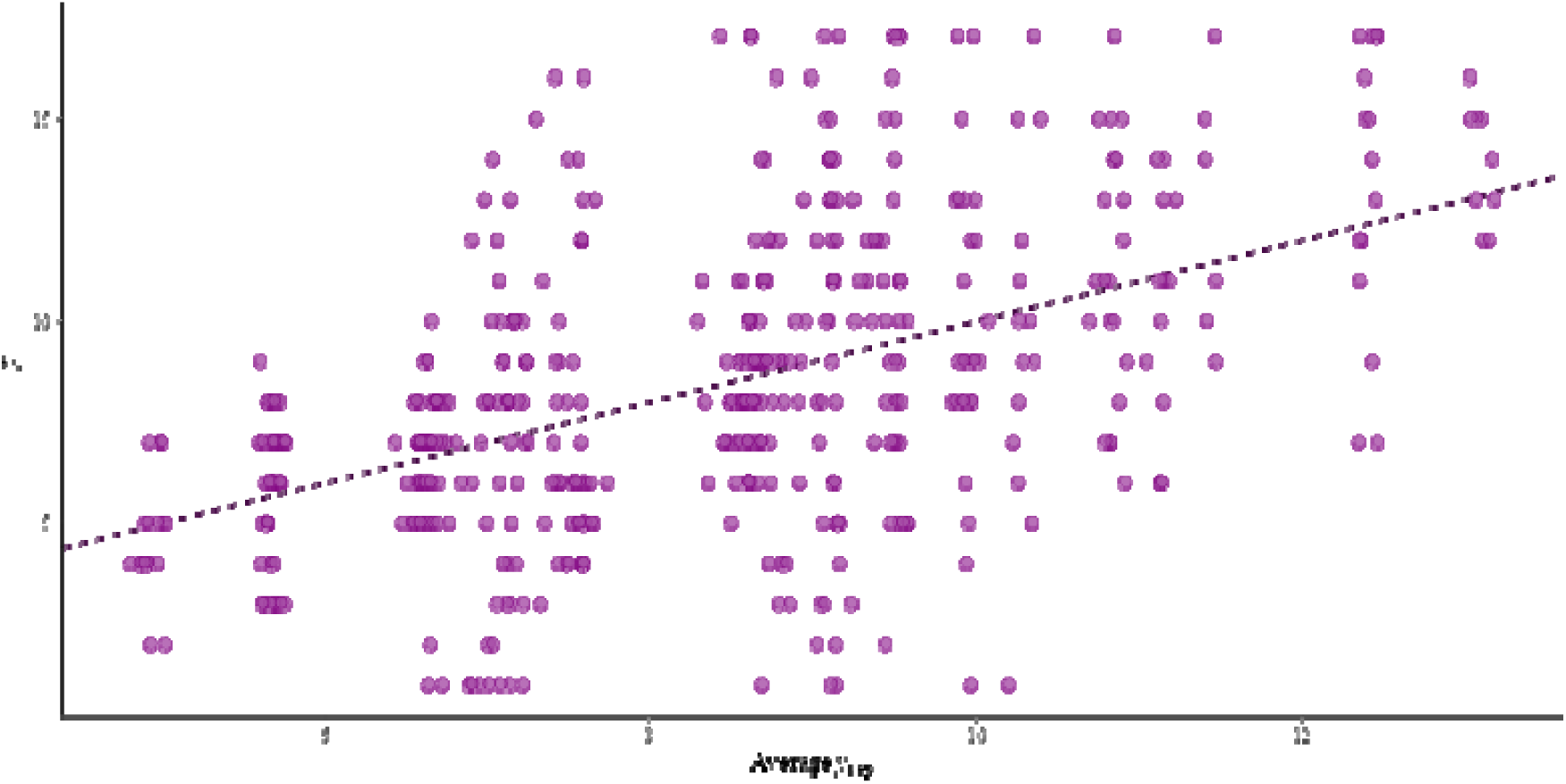
Observations (y) vs average simulated values (y_rep_).

**Figure S5:**
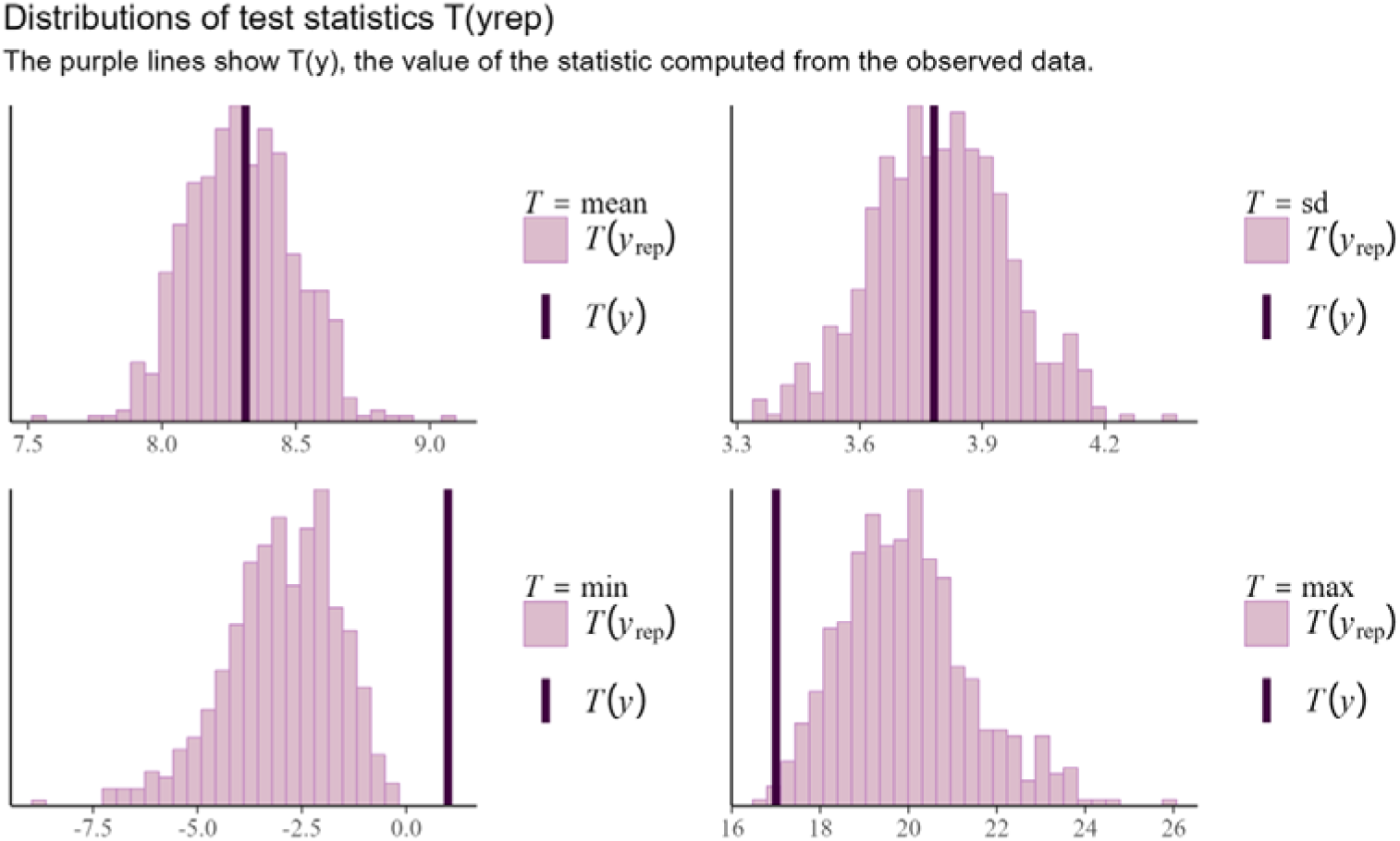
Distributions of test statistics *T*(*y_rep_*) compared to the value of the statistic computed from the observed data T(y) (the dark purple lines).

**Figure S6:**
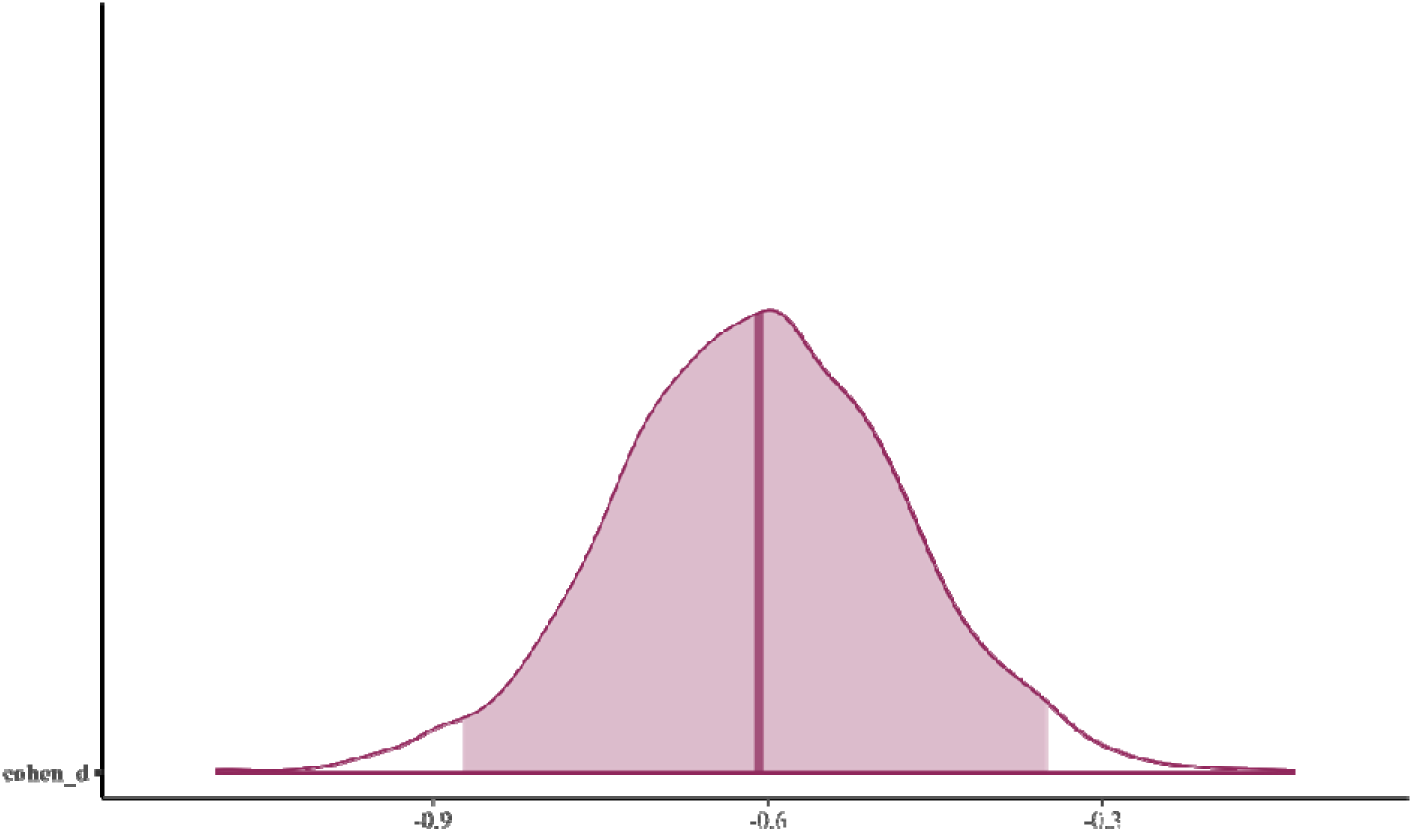
Cohen’s d, indicating a medium effect size estimate for the observed difference in phenology timing between natural and restored meadows. The value is calculated by dividing the difference between the two group means by the data’s standard deviation.

